# An enzyme activation network provides evidence for extensive regulatory crosstalk between metabolic pathways

**DOI:** 10.1101/2023.11.16.567372

**Authors:** Sultana Mohammed Al Zubaidi, Muhammad Ibtisam Nasar, Markus Ralser, Richard A. Notebaart, Mohammad Tauqeer Alam

## Abstract

Enzyme activation by cellular metabolites plays a pivotal role in regulating metabolic processes. Nevertheless, our comprehension of such activation events on a global network scale remains incomplete. In this study, we conducted a comprehensive investigation into the optimization of cell-intrinsic activation interactions within *Saccharomyces cerevisiae*. To achieve this, we integrated a genome-scale metabolic model with enzyme kinetic data sourced from the BRENDA database. Our objective was to map the distribution of enzyme activators throughout the cellular network. Our findings indicate that virtually all biochemical pathways encompass enzyme activators, frequently originating from disparate pathways, thus revealing extensive regulatory crosstalk between metabolic pathways. Indeed, activators have short pathway lengths, indicating they are activated quickly upon nutrient shifts, and in most instances, these activators target key enzymatic reactions to facilitate downstream metabolic processes. Interestingly, non-essential enzymes exhibit a significantly higher degree of activation compared to their essential counterparts. This observation suggests that cells employ enzyme activators to finely regulate secondary metabolic pathways that are only required under specific conditions. Conversely, the activator metabolites themselves are more likely to be essential components, and their activation levels surpass those of non-essential activators. In summary, our study unveils the widespread importance of enzymatic activators, and suggests that feed-forward activation of conditional metabolic pathways through essential metabolites mediates metabolic plasticity.

## Introduction

The metabolic network, the largest interconnected system of the cell ^1–3^, is an essential biological system by which organisms produce necessary components such as energy molecules and cellular building blocks including carbohydrates, lipids, nucleotides, proteins, and vitamins for their growth and survival ^3^. To survive in changing environments, the regulation of cellular compounds is required ^4^. Cells achieve metabolic regulation by a range of regulatory interactions ^5,6^. The most common mechanisms of regulatory interactions are occurring at different levels of cellular processes. These include hierarchical control, which includes transcriptional and translational level regulations to change the enzyme level, the posttranslational modification of enzymes to change their activity, and metabolite-enzyme interactions that can inhibit or activate enzymes. ^7–9^ As the latter is driven by cell intrinsic metabolites that are formed within or outside the regulated pathway, the metabolic network is in part self-regulatory ^10–12^. Characterizing these metabolite-enzyme interactions is thus key to understanding metabolic phenotypes.

Broadly speaking, enzyme inhibitory regulations can be allosteric, non-competitive and competitive. Among these, most frequent are the competitive interactions, in which the inhibitor competes with the substrates to bind to the enzyme’s active site ^13^ reducing the rate of enzymatic reactions ^14^. Competitive inhibitors are often the by-product of the same biochemical pathway, or created in close proximity within the metabolic network ^13^. Indeed, enzyme inhibitory interactions in human metabolism emerge merely due to a finite chemical diversity that prevails within the metabolome ^13^.

In contrast, enzyme activation interactions enhance the rate of metabolic reactions, and most activators bind allosterically ^15^. For example, Phosphofructokinase-1 (PFK-1, EC: 2.7.1.11), one of the most important glycolysis regulatory enzymes, is allosterically activated by a number of cellular metabolites including ADP, GDP, AMP, PI, Glutathione, but the most potent activator of PFK-1 is fructose 2,6-bisphosphate (F-2,6-BP) that is produced by Phosphofructokinase-2 enzyme (PFK-2, EC: 2.7.1.11). While individual examples have been well studied, the extent of activation interactions is not entirely characterized at a cellular-wide level.

In this study, we have reconstructed a network of enzyme-metabolite activatory interactions by combining the genome-scale metabolic network of *Saccharomyces cerevisiae* ^16^ with enzyme activation data from the Braunschweig Enzyme Database (BRENDA) database, which has collected enzyme kinetic data over a century of biochemical research ^17^. The cell-intrinsic enzyme-metabolite activation network shows that up to 40% of enzymatic reactions are intracellularly activated. Activated enzymes are found across the entire metabolic system, covering most of the biochemical pathways. We show that while most of the highly activated enzymes are non-essential, the highly activating compounds are essential for growth and are often produced shortly after nutrient uptake. Further, we have compared the enzyme-metabolite activation interaction with the genetic interaction and examined the co-expression pattern of the genes associated with activating metabolites and activated enzymes by analyzing transcriptome profiles from more than 600 unique conditions from >200 experiments ^18^, and conclude that the cell-intrinsic metabolic activation interactions could be used to understand the observed metabolic profile in various conditions.

## Results

### Cell-intrinsic activation network of *Saccharomyces cerevisiae*

We have reconstructed a genome-scale enzyme-metabolite activation interactions network of *Saccharomyces cerevisiae.* For each metabolic enzyme (in total 469 enzymes) in the genome-scale metabolic model (iMM904) ^16^ a list of all associated activatory molecules were downloaded from the Brenda database, using SOAP clients instructions ^17^. The obtained activators list contains all kinds of molecules including both non-cellular molecules as well as intracellular metabolites, all increasing the enzyme’s activity as determined with enzyme assays. By comparing the list of activatory molecules with the intracellular metabolites of the metabolic model ^16^ all non-cellular molecules were removed (Figure 1a; see the method section for details) to produce the cell-intrinsic activation interactions network (Figure 1b, Supplementary File 1). In the network, the nodes represent enzymes and activator metabolites, and the edges are formed between nodes when an enzyme is activated by a metabolite (Figure 1b). The final network consists of 874 activatory interactions associated with 187 enzymes and 201 cellular metabolites, and like most other biological networks, the cell-intrinsic activation interactions network is scale free and follows the power law distribution (Supplementary Figure S1).

**Figure 1:**
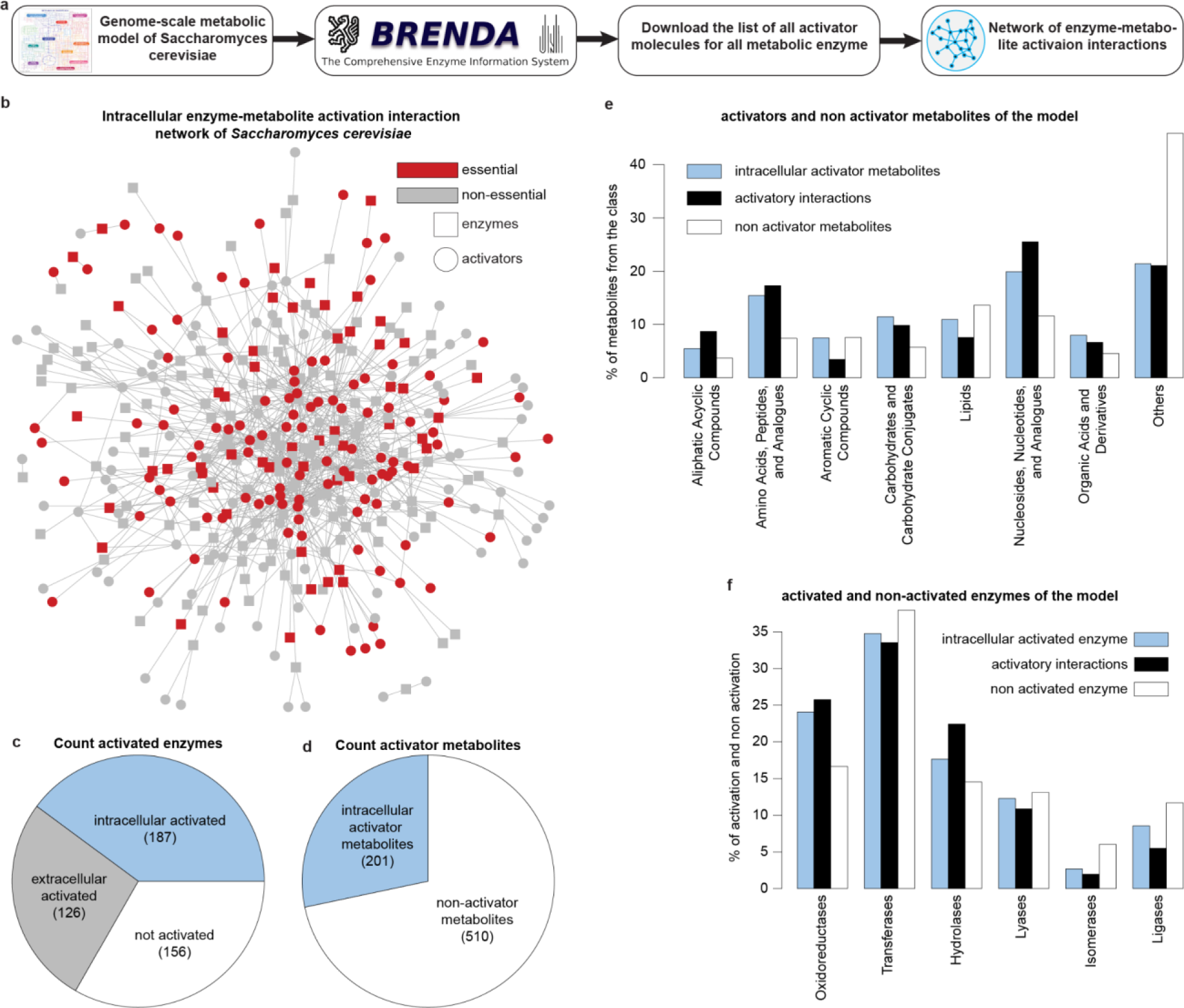
cell-intrinsic enzyme-metabolite activation interactions network. (a) Schematic of reconstructing cell-intrinsic enzyme-metabolite activation interactions network. (b) The network of enzyme-metabolite activation interactions within Saccharomyces cerevisiae contains 187 enzymes and 201 metabolites as nodes, and they are connected by 874 links representing enzyme-metabolite activation. The essential and non-essential components are marked in red and gray, respectively. (c) Analysis of the distribution of enzymes across metabolism, categorizing them as intracellularly activated, extracellularly activated, or unactivated. (d) Proportion of activator and non-activator metabolites within the metabolic network. (e) The prevalence of intracellular activator metabolites, non activator metabolites, and activatory interactions within different compound classes. (f) The prevalence of intracellular activated enzymes, activatory interactions, and non activated enzymes within different enzyme classes.

In terms of the metabolic network coverage, 40% of the total metabolic enzymes (187 out of 469) are intracellularly activated, and the remaining 60% of the metabolic enzymes (282 out of 469) are either activated by extracellular molecules (126; 27%) or contain no activation interactions at all (156; 33%) (Figure 1c). On the other hand, out of the total 710 metabolites, 28.3% (201 in total) act as activators on at least one enzyme (Figure 1d). To understand the prevalence of activators within different compound classes, all cellular metabolites were clustered into 8 different compound groups based on the Kegg database classification ^19^ (Figure 1e). Lipids, which are highly common within inhibitory metabolites ^13^, have the low prevalence of activatory metabolites, and are associated with even fewer activatory interactions (Figure 1e). On the other hand, metabolites belonging to ‘Nucleosides, Nucleotides, and Analogues’, ‘Amino Acids, Peptides, and Analogues’, ‘Carbohydrates and Carbohydrate Conjugates’, ‘Aliphatic Acyclic Compounds’, and ‘Organic Acids and Derivatives’ have substantially higher prevalence of activatory metabolites compared to non activator metabolites (Figure 1e). Besides, the maximum percentage of activatory interactions are conducted by the ‘Nucleosides, Nucleotides, and Analogues’ (∼25%), which explains the important roles of this metabolite group in positively regulating metabolic reactions (Figure 1e).

In the network of human enzyme inhibition interactions ^13^ all enzyme classes were equally susceptible to metabolic inhibitions. In contrast, in the activation network (Figure 1b) each enzyme class has a different prevalence of activated enzymes (Figure 1f). Out of the total intracellular activated enzymes, almost 35% belong to Transferases that is also the largest enzyme class of the metabolic network ^16^, and are equally associated with all cell-intrinsic activated interactions (34%) (Figure 1f). Isomerases, and Ligases, which are the two smallest enzyme classes of the metabolic network, have even lower ratios of activated enzymes compared to non-activated enzymes (2.6%, and 8.5% respectively) (Figure 1f). In contrast, Oxidoreductases and Hydrolases, which are 2nd and 3rd largest metabolic enzyme classes, catalyzing thermodynamically favored metabolic reactions (non-equilibrium), have substantially higher ratios of intracellularly activated enzymes (24% and 17.6% respectively, Figure 1f).

### Activatory interactions are distributed across metabolism, regulating thermodynamically favorable reactions

A substantially low percentage of intracellularly activated enzymes (40%, Figure 1c) spans throughout the metabolism, covering the majority of the metabolic pathways that have more than 2 enzymatic reactions (59 out of 60; 98.33%) (Figure 2a, b). Some of the highly intracellularly activated metabolic pathways are Pyruvate metabolism (14/19), Glycolysis / Gluconeogenesis (13/20), Purine and Pyrimidine metabolism (12/34 and 9/19 respectively), and various amino acid metabolic pathways such as Alanine, aspartate and glutamate metabolism, Glycine, serine and threonine metabolism Cysteine and methionine metabolism, Arginine biosynthesis, Phenylalanine, tyrosine and tryptophan metabolism, and Tryptophan metabolism (Figure 2b). We also observe that some of the metabolic pathways are enriched with only essential activated enzymes (for instance, Phenylalanine, tyrosine and tryptophan biosynthesis, Terpenoid backbone biosynthesis, and Fatty acid biosynthesis) whereas in some pathways none of the activated enzyme seems essential for growth (for example Glycolysis / Gluconeogenesis, Citrate cycle (TCA cycle), Galactose metabolism, Tryptophan metabolism, and Cyanoamino acid metabolism).

**Figure 2:**
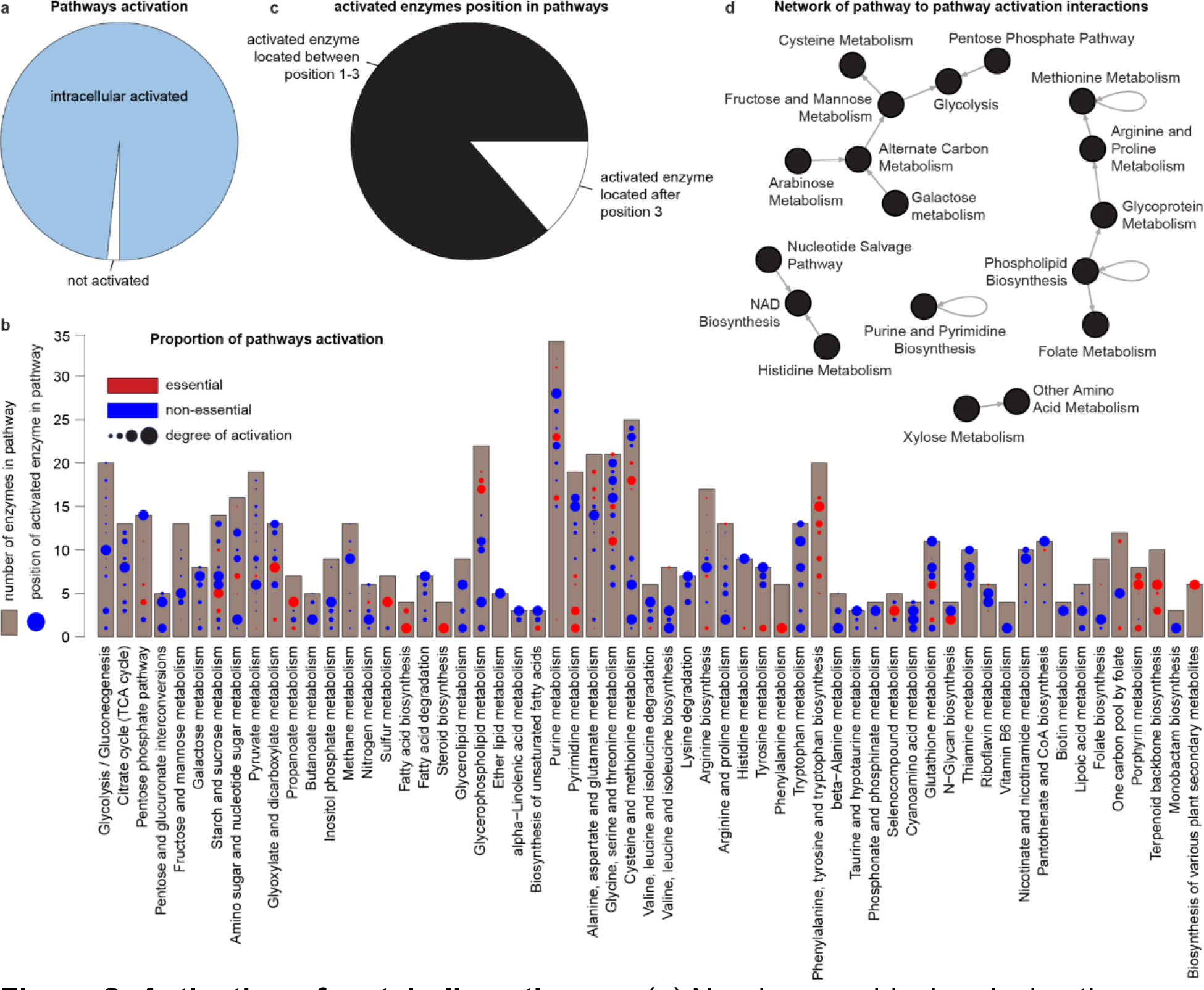
Activation of metabolic pathways. (a) Nearly every biochemical pathway exhibits activatory interactions (59 out of 60). (b) The activated enzymes, either essential or non-essential with varying degrees of activation, are distributed across metabolic pathways. (c) Remarkably, most of the biochemical pathways (86.4%) include at least one enzymatic reaction in the initial three positions of the pathway. (d) The interconnected network of pathway to pathway activation highlights trans-activation wherein metabolites from one pathway notably stimulate enzymes in other pathways. This stands in stark contrast to the inhibitory interactions network described by Alam et al. in 2016, where the majority of pathways primarily inhibit their own enzymes. The enzyme localization in metabolic pathways is according to KEGG pathways definition.

Furthermore, we examined the position of these activated enzymes in metabolic pathways. We find that at least one of the first three initial enzymatic reactions of most of the pathways (51/59; 86.4%) are positively regulated (Figure 2c). In biochemical pathways, reactions at the beginning are generally non-equilibrium flux-generating reactions which must be regulated to change the flux through the entire pathway ^20^. Additionally, we also observed that the activated enzymes are not only restricted to the beginning of the biochemical pathways rather they are distributed throughout the pathways. This spread of activated enzymes shows that the other distant reactions could also be regulated by intracellular metabolites depending on conditions. However, on the other hand, we observe that out of 122 enzymes, which are located after position 3 in various chemical pathways, 38 enzymes (31%) are also placed between position 1-3 in other pathways. Further, it is important to note that the average length of pathways (number of enzymes) is 10.2, 37% of the pathways have more than 10 enzymes, and 73% pathways have more than 5 enzymes. Lastly, we did not observe any particular pattern of enzyme essentiality or the degree of activation for early or late regulated enzymes of the pathway (Figure 2b).

### Evidence for a high degree of trans-activation within the metabolic network

Next, we investigated how different biochemical pathways significantly activate each other (Figure 2d). In many pathways, we detect feed-forward and feedback activation, for instance, methionine metabolism, purine and pyrimidine biosynthesis, and phospholipid biosynthesis. However, our network provides evidence for a significant degree of trans activation between pathways. For instance, glycolysis is significantly activated by fructose and mannose metabolism, and pentose phosphate pathway. Moreover, its products are insignificantly activating any other metabolic pathways. This highlights the importance of intracellular activation of glycolysis which occurs at the beginning of the metabolism to feed the rest of the metabolic network. Similarly, NAD biosynthesis is activated by two different pathways including histidine metabolism, and nucleotide salvage pathway. Out of the 18 central metabolic pathways from the network of pathway-to-pathway activation interactions, 6 pathways including Cysteine Metabolism, Other Amino Acid Metabolism, Folate Metabolism, Glycolysis, NAD Biosynthesis, and Purine and Pyrimidine Biosynthesis are only activated by other pathways. Except for self activation in 3 pathways, they do not activate other pathways. Moreover, 7 pathways including Histidine Metabolism, Nucleotide Salvage Pathway, Xylose Metabolism, Arabinose Metabolism, Galactose metabolism, Phospholipid Biosynthesis, and Pentose Phosphate Pathway are sources of activation for other pathways and they are not activated by other different pathways. Indeed, several central metabolic pathways (15/18) are significantly activating or activated by other pathways (Figure 2d).

### Highly activating metabolites are mostly essential whereas highly activated enzymes are non essential for growth carrying less gene expression variation

The essentiality of enzymes and metabolites, predicted using the flux balance analysis approach for an optimal growth, was combined with the cell-intrinsic activation network (Figure 1b). We observe a significant difference in the prevalence of essentiality of activators compared to non-activator metabolites (P = 2.59e-20, Fisher’s Exact Test). More than half of the cellular activator metabolites are essential (53.23%) for growth, whereas the essentiality ratio is merely 20% for non activators (Figure 3a). In contrast, activated and non-activated enzymes have no significant difference in the prevalence of essential enzymes (P = 0.97, Fisher’s Exact Test), although non-activated enzymes are slightly more prevalent with essential enzymes (36.8%) compared to activated enzymes (28.8%) (Figure 3a).

**Figure 3:**
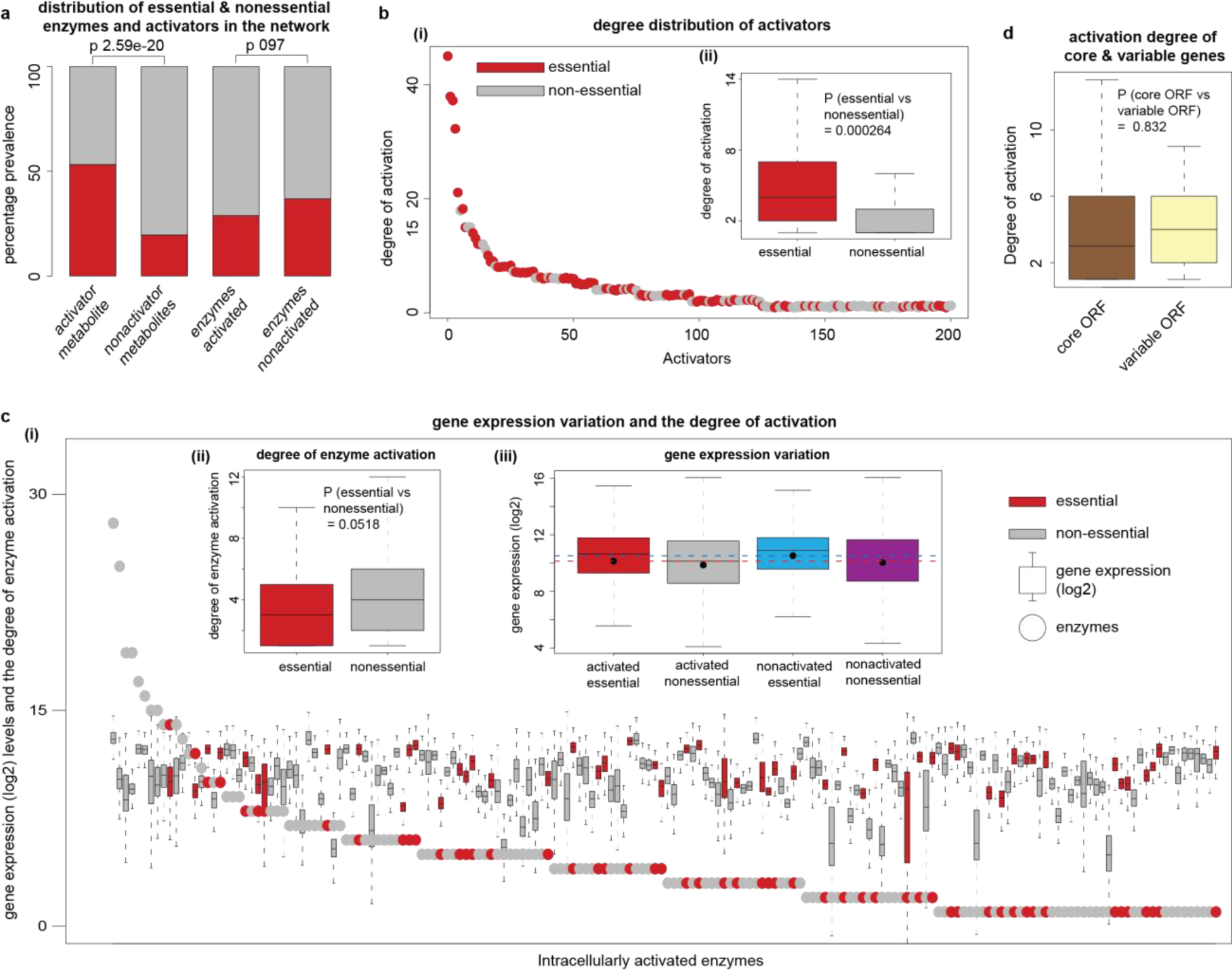
Degree distribution of activators and enzymes. (a) In contrast to non-activator metabolites, activators demonstrate a notable increase in essential metabolites (p = 2.59e-20). However, when it comes to enzymes, there are no significant differences in essentiality levels between activated and non-activated enzymes (p = 0.97). (b) The degree distribution of activator metabolites within the cell-intrinsic activation interaction network reveals that highly interactive activators are mostly essential for growth (i). Furthermore, metabolites essential for growth show significantly more activatory interactions compared to non-essential activatory metabolites (P = 0.000264) (ii).(c) Conversely, the distribution of degrees among activated enzymes in the cell-intrinsic activation interaction network indicates that highly activated enzymes are not essential. Moreover, the variability in gene expression data across over 500 conditions for all activated enzymes, whether essential or non-essential, was assessed in relation to the degree of interactions. (i). Strikingly, essential enzymes have a significantly lower degree of activation compared to non-essential enzymes (P = 0.05) (ii). Moreover, variations are evident in the global gene expression profiles among activated essential enzymes, activated non-essential enzymes, non-activated essential enzymes, and non-activated non-essential enzymes (iii). (d) Finally, a comparison was conducted regarding the degree of activation between core genes and variable genes, derived from xyz et al . Core genes, characterized by conservation across strains, exhibit a generally lower average number of activation interactions compared to variable genes, which lack conservation across species.

We analyzed the degree of activation for both essential and nonessential metabolites, and found that 6 out of 7 hub activator metabolites, activating more than 15 enzymes, are essential (Figure 3b i). In concurrence, the number of interactions associated with essential metabolites is significantly higher compared to non-essential activators (P = 0.000264, T test), where 53% of the essential activatory metabolites are activating more than 70% of the interactions in the network (Figure 3b ii).

In contrast, the highly activated enzymes seem to be non-essential for growth, where the degree of activation for non-essential enzymes is significantly higher compared to the essential enzymes (P = 0.05, T test) (Figure 3c i, ii). We find all of the hub enzymes, which are activated by more than 15 metabolites (degree>15), are non-essential for growth (Figure 3c i). It has previously been reported that essential enzymes are tightly regulated in order to maintain the constant level of gene expression irrespective of the genetic or environmental perturbations ^11,21^, therefore, such genes are expected to have less variation in the gene expression ^22^. In concordance with other studies, we find that the essential enzymes have higher gene expression levels across conditions compared to non-essential enzymes(average log2 gene expression 10.3 vs 9.87). We also observe that the activated enzymes have slightly lower average gene expression levels compared to non-activated enzymes (average log2 gene expression 10.01 vs 10.19). Enzymes which are essential for growth and independent from any activation interactions have on average higher gene expression values and less variance (average log2 gene expression 10.53, SD 1.72, Figure 3c iii). In contrast, the enzymes which are not essential for growth and dependent on activation for their expression, possess the lowest average gene expression across conditions and highest variance (average log2 gene expression 9.87, SD 2.18, Figure 3c iii). Furthermore, we find that the degree of enzyme activation associated with core genes, which are conserved across all *S. cerevisiae strains*, is slightly lower compared to variable genes, which are not conserved across strains (Figure 3d). This further explains that the highly conserved and essential enzymes are generally independent from required activation.

### Activatory metabolites are produced shortly after nutrient uptake

We have analyzed the metabolic network topology and examined how activator and non activator metabolites are produced within the metabolic network. First, we converted the metabolic model ^16^ into a graph where nodes represent metabolites and edges are established when two metabolites are connected by a chemical reaction. Using this network, we calculated the shortest path length, representing the minimum number of chemical transformations required to produce different activators and non-activator metabolites from glucose metabolite, the main carbon source nutrient. We find that within the metabolic network the shortest path length from glucose to activators is significantly lower compared to non-activators metabolites (Figure 4a, P = 1.52e−17 (t test)). The majority of the activator metabolites have the shortest path length of 5 whereas for non-activator this distance ranges from 5-7 (maximum length 6, Figure 4b). In addition, we also observe a strong correlation between shortest path length and the degree of activation for activating metabolites, where the activators with the lowest shortest path length (>=5) seem to activate most of the enzymatic reactions (Figure 4c). For instance, activators with the shortest path length of 5 are activating up to 40 enzymatic reactions whereas four metabolites which have the shortest path length of 7 from glucose metabolite are activating at most only 5 reactions (Figure 4c), and a metabolite with shortest path length 9 activates only one enzyme. Furthermore, we also find a significant difference between the shortest path length considering all exchange nutrient metabolites to activators compared to non-activators (Supplementary Figure S2). These results suggest an inbuilt optimality principle for producing cellular activator metabolites within the metabolic network. This highlights that intracellular super activators are produced shortly after nutrient uptake which could be later utilized for positively driving the overall metabolism. Similar to this observation, other studies have reported a well-characterized set of 12 metabolites, which are precursors for all essential biomass components of the cell. These precursor metabolites, located at different branchpoint nodes, are converted from carbon source nutrients by shortest path length, suggesting an optimality principle for central carbon metabolism.

**Figure 4:**
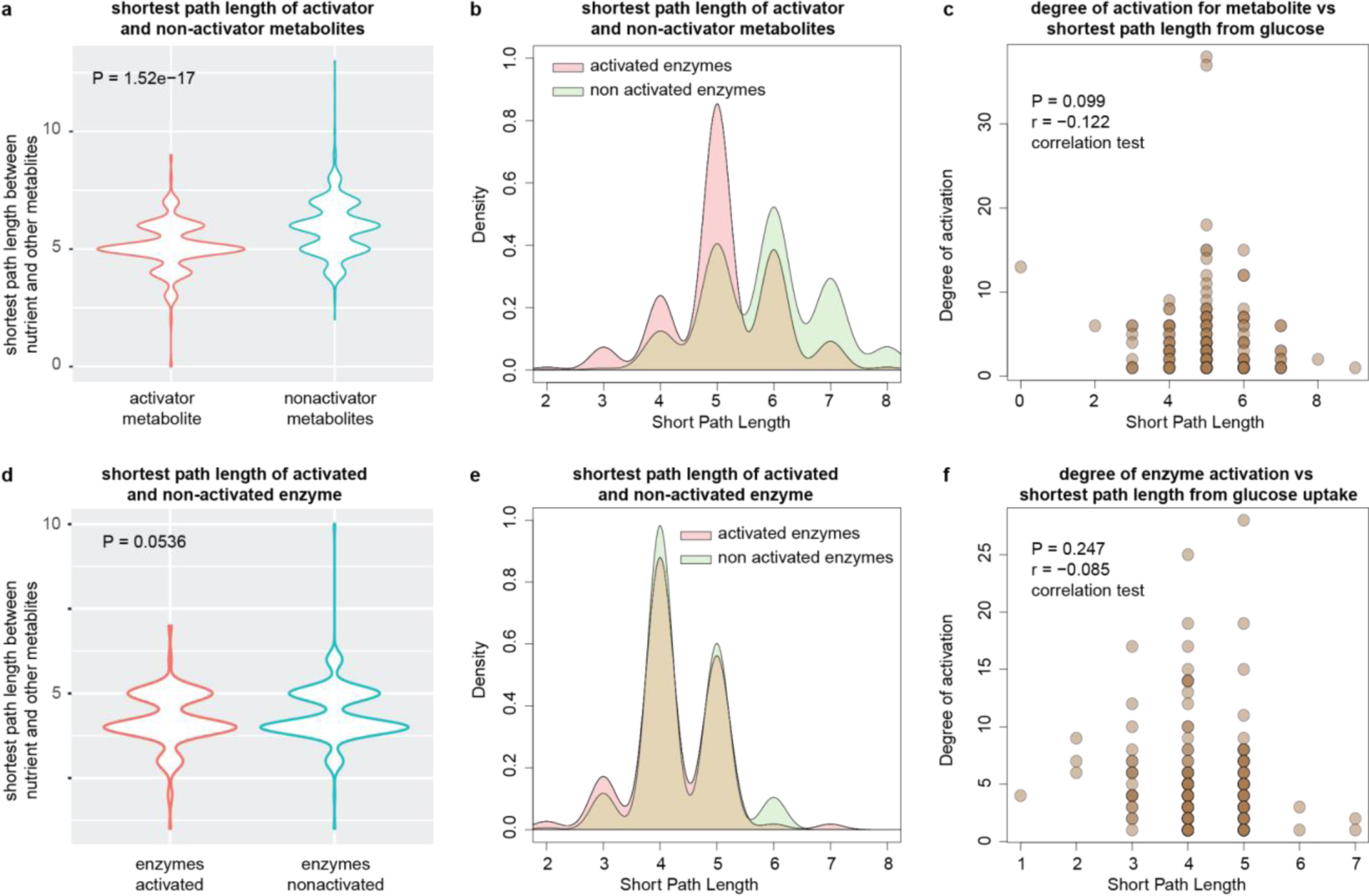
Shortest path of activator metabolites and activated enzymes in the metabolic network. (a) Within the organism’s metabolic network, the shortest path length was computed for all metabolites from glucose, the reference nutrient metabolite. Activator metabolites exhibit a significantly shorter shortest path length compared to non-activators (P = 5.77e−36, t-test). (b) Additionally, a majority of activators have a shortest path length of 5, while for non-activators, it is 6. (c) Analyzing the correlation between the shortest path length and the degree of activation reveals a negative association between the two. (d) Likewise, the shortest path length was determined for all enzymes originating from the glucose transporter, serving as the reference nutrient uptake enzyme. While a statistically significant distinction exists between activated and non-activated enzymes concerning their shortest path length from the glucose transporter (P = 0.05), (d) this disparity is primarily attributed to a minimal fraction of non-activated enzymes with a slightly higher shortest path length (that is 6). Beyond this, no variation is observed in the shortest path length between activated and non-activated enzymes. (e) Finally, there is no discernible correlation between the degree of enzyme activation and their respective shortest path lengths.

In contrast, we did not observe any major difference between activated and non-activated enzymes in terms of the shortest path length, considering glucose exchange reaction as the root (Figure 4d, e). For the majority of activated and non-activated enzymes, the shortest path length is 4-5 (Figure 4d, e). Furthermore, we find that the enzymes with shortest path length of 4-5 have substantially higher degree of activation compared to enzymes with shortest path length of 6-7 (Figure 4f). This is also due to the fact that most enzymatic reactions have the shortest path length of 4-5 and very few reactions are distant from the glucose transport reaction.

### Coexpression of genes associated with activatory interactions

We have analyzed the co-expression pattern of genes linked with nodes of every interaction of the cell-intrinsic activation network. For this analysis, highly connected hub metabolites such as Phosphate, AMP, ADP and ATP molecules were removed due to their connection with hundreds of genes. Genes linked to the metabolic reaction for producing activatory metabolites and activated enzymes were based on the metabolic model definition (iMM904). The gene expression data from 207 experiments consisting of 665 unique conditions were obtained from the ArrayExpress database (microarray experimental data from Genome2 chips), and processed using the limma package in R to produce the gene expression profile for every gene (please see the method section for details).

For each pair of genes, corresponding to the nodes of the cell-intrinsic activation interaction, we have performed the correlation test, and considered the genes co-expressed based on a stringent threshold cutoff (r>0.75 and P adjusted <0.001). Figure 4a shows a subnetwork of the main cell-intrinsic activation interaction in which genes associated with both nodes are co-expressed. Out of 758 cellular activatory interactions (after removing cofactors from the main network), only 211 interactions (28%), produced from 69 metabolites and 83 enzymes, have coexpressed gene expression across conditions (Fig 5a, b), which highlight that such interactions are quite common in constraining the metabolism in different conditions. Further, by comparing activatory interactions with genetic interaction networks we find that 33% of the interactions are also genetically interacting, where genes associated with such interactions are genetically linked. Out of the total co-expressed activatory interactions, 59 enzymes, corresponding to 157 interactions, are non essential and only 24 enzymes are essential for growth (Fig 5c). However, for the activators we find 48 metabolites are essential (Fig 5d).

**Figure 5:**
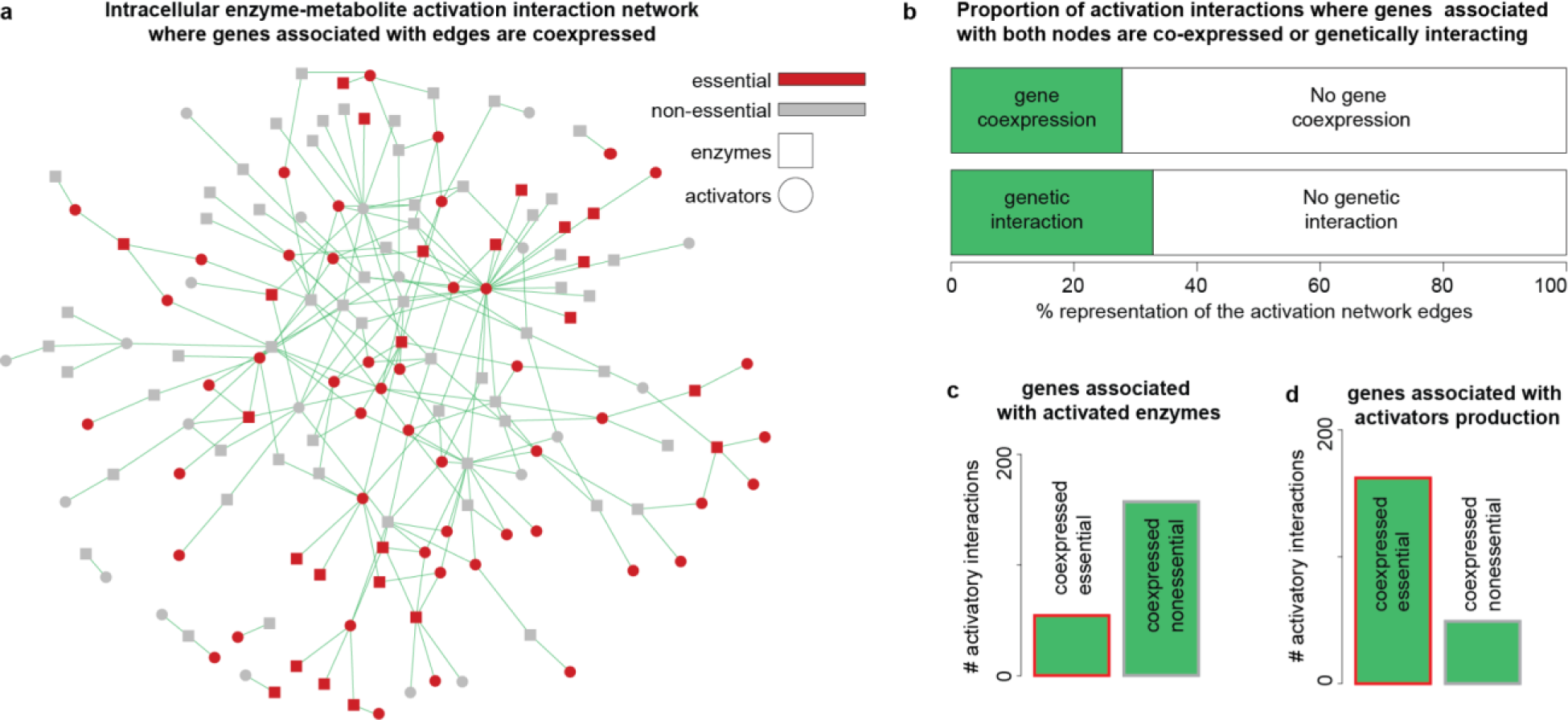
The global co-expression profile of genes associated with activatory interactions. (a) A portion of the activation network exhibiting co-expression of genes linked to both enzyme and metabolite nodes in the connection. The subnetwork highlights both essential and non-essential metabolites and enzymes. (b) Within the activation network, approximately 27.8% of the total connections display notable co-expression between genes linked to activator production and the activated enzymes, as evidenced by a correlation coefficient r > 0.7 and P < 0.01. Similarly, concerning the global genetic interactions in S. cerevisiae, nearly 35% of interactions involve both essential genes. (c) The distribution of enzymes categorized as co-expressed and essential is notably lower than that of genes co-expressed and non-essential. (d) Conversely, for genes associated with activator metabolites, the count of co-expressed and essential genes is significantly higher than that of co-expressed and non-essential genes.

## Discussion

The metabolic network possesses inherent regulatory mechanisms to autonomously control its functionality by modulating enzyme activity through various enzyme-metabolite interactions ^23–26^. These interactions can either be inhibitory, where the metabolite diminishes the enzyme’s activity, or activatory, where cellular metabolites enhance the enzyme’s activity ^2,13,24,27,28^. In the metabolic system, numerous enzyme-metabolite inhibitory interactions exist, acting as constraints on metabolism. Studies indicate that over 75% of inhibitory interactions in humans are competitive, wherein structurally similar substrates and metabolites compete for the same active site. Consequently, inhibitory metabolites are frequently produced within the same pathway and significantly impede their own enzymatic reactions ^13^. However, comprehensive global characterization of activation interactions remains incomplete.

In this study, we systematically examined the cell-intrinsic activation interactions on a global scale for *Saccharomyces cerevisiae*. Unlike most inhibitory interactions, the enhancement of enzyme activity in this context necessitates allosteric binding of metabolites, facilitating the substrate binding to the active site ^29^. To comprehensively grasp the global activation interactions within the metabolic system, we reconstructed the network of enzyme-metabolite activation interactions. This involved processing activator compounds from the BRENDA database ^17^ and mapping them with the genome-scale model of *Saccharomyces cerevisiae* (iMM904) ^16^. The cellular intrinsic activation network comprises 187 enzymes and 201 metabolites, interconnected by 874 activatory interactions (Fig 1b). It is important to note that our approach operates under the fundamental assumption that metabolic activators are conserved ^30,31^. However, a limitation of our methodology is the lack of adequate data in the BRENDA database to exclusively construct the yeast-specific activator network based on yeast enzymology data. However, not all enzyme activators will be conserved ^30,32^, and thus we might overestimate the number of activators for some reactions.

Initially, we investigated the extent to which cellular metabolites activate enzymatic metabolic reactions within the metabolic system and their distribution across various biological processes. Our findings reveal that 40% of the total metabolic enzymes are activated by cellular metabolites (Fig 1c). Among the remaining 60% of enzymes, approximately half are reported to be activated by external metabolites, while the rest show no activation (Fig 1c). Notably, the activated enzymes are distributed throughout the metabolic network, encompassing a majority of biochemical pathways (Fig 2a). This comprehensive coverage of metabolic pathways with intracellularly activated enzymes elucidates the optimality of inherent cell-intrinsic activation constraints on metabolism, underscoring their crucial role in the efficient operation of metabolic pathways ^5^. Additionally, prior studies employing metabolic control analysis and flux coupling analysis support the observation that metabolic controls are widely dispersed across the metabolic network ^10,33^.

Moreover, in the majority of biochemical pathways, at least one of the top three enzymatic reactions is activated intracellularly (**Fig 2 b, c**). This underscores the remarkable inherent mechanism within the cell for effectively controlling these initial reactions, thereby regulating the entire metabolic network efficiently. Since these initial reactions primarily involve flux-generating non-equilibrium reactions ^34–36^, they are frequently regulated to facilitate pathway flux ^37^. The intracellular activation of key enzymatic reactions in most biochemical pathways is a response to the imperative requirement for precise regulation and control of cellular metabolic processes ^5,6,28,33^. Interestingly, we observe activated enzymes at various positions in all biochemical pathways, even though the activation of individual enzymatic reactions might not consistently benefit the entire pathway (Fig 2b). One rationale for this finding is that around one-third of enzymatic reactions found in the middle or later stages of a pathway are also commonly present at the start of other biochemical pathways (Fig 2b). Activating the remaining non-initial enzymatic reactions could be essential for maintaining flux through the pathway after the initial reactions are activated ^36,38^. Moreover, the presence of activated enzymes at different positions in biochemical pathways underscores the complex and adaptable nature of cellular metabolism ^36^. This configuration empowers cells to meticulously regulate their metabolic activities, respond to diverse signals, and optimize their functionality in various scenarios ^36,39,40^.

We further explored the extent to which metabolites from one pathway significantly activate enzymes in other pathways (Fig 2d). This investigation is prompted by the prevailing allosteric binding of activators with enzymes, facilitating the binding of substrates to the enzyme ^29^. It is noteworthy that activator metabolites and substrates exhibit structural dissimilarity owing to this widespread allosteric binding mechanism. Additionally, prior research has illustrated that metabolites within the same pathways demonstrate greater structural similarity in comparison to those across different pathways, often leading to their association with competitive binding with enzymes—the most common inhibitory interactions ^13^. Interestingly, the configuration of the activation interactions network between pathways stands in stark contrast to that of the human inhibitory interactions network. In the latter, products originating from the same pathway often act as significant inhibitors ^13^, whereas the activatory interactions between pathways are mostly cross-pathway in nature (Fig 2d). This disparity underscores the distinct nature of activation and inhibition interactions within the metabolic network, playing a crucial role in constraining metabolism for optimal efficiency.

In terms of activation of different enzyme classes, transferases, oxidoreductases, and hydrolases, which are also the most prevalent enzyme classes in the metabolic network in that order, cover most of the activated enzymes. In particular, hydrolases and oxidoreductases have a higher prevalence of activated enzymes compared to non-activated enzymes (Fig 1f), which could be due the nature of their catalytic activities and the functional requirements of these enzyme classes. Hydrolases, for instance, catalyze the hydrolysis of various substrates by adding water molecules ^41^. Activators can modulate the enzyme’s conformation, making it more receptive to substrate binding and efficient hydrolysis. Similarly, oxidoreductases are involved in oxidation-reduction reactions, fundamental to cellular processes, where electrons are transferred between substrates ^42^. Activation of these enzymes by specific metabolites allows for precise control of electron transfer and redox balance in response to cellular needs. In contrast, isomerases and ligases, which are the least prevalent enzymes in the metabolic network, have an even lower percentage of activated enzymes (Fig 1f). These enzyme classes typically engage in enzymatic activities that are inherently less complex or less dependent on allosteric regulation. Isomerases facilitate structural rearrangements, which may not require extensive external modulation. Ligases, on the other hand, often have an inherent regulatory mechanism through the energy input required for bond formation ^43^. This over or under representation of intracellular activation highlights the importance of activation for specific classes of enzymatic reactions.

Further, within the list of activated enzymes, we find 28.8% of enzymes are essential for growth in a minimal media condition; a similar proportion of essentiality was also observed for non-activated enzymes (36.8%) (Fig 3a). Strikingly, the essential activated enzymes have significantly less degree of activation compared to non-essential enzymes (Fig 3c). This includes the topmost activated enzymes such as EC:2.7.1.40, EC:1.4.1.2, EC:3.1.3.2, EC:3.1.3.5, EC:2.7.1.11, and EC:3.5.1.2 which are intracellularly activated by 28, 25, 19, 19, 17 and 17 different metabolites, respectively, which are non-essential for growth. This remarkable significant difference in the degree of activation between essential and nonessential enzymes may be a result of the evolutionary and functional characteristics of these enzymes ^36,39^. The role and involvement of essential enzymes in core cellular processes might necessitate a more streamlined and efficient mode of regulation, with less reliance on activator interactions.

We then examined which cellular metabolites are optimal to act as an activator. For cellular activator metabolites, we find strikingly opposite features compared to activated enzymes (Fig 3). We observe that the ratio of essential metabolites among the activators is significantly higher compared to non-activator metabolites (53% essential activators vs 25% essential non-activators) (Fig 3b). In addition, the essential metabolites activate a significantly higher number of enzymatic reactions compared to non-essential metabolites (Fig 3b, Fig 1e). Some of the highly activating metabolites are C00002, C00051, C00097, C00009, and C00020 activating 45, 38, 37, 32, and 21 different enzymes, respectively. Since essential metabolites are consistently generated within the cell, regardless of the conditions, employing them to also activate other enzymes might be a strategic and resource-efficient approach ^44^ Furthermore, the conservation of crucial metabolic pathways could also contribute to the predominance of essential metabolites serving as activators.

Lastly, we computed the shortest path length for all cellular metabolites from different nutrient sources, including glucose. Our findings indicate that activators exhibit significantly shorter path lengths compared to non-activators (Fig 4). This suggests that activator metabolites are promptly produced following nutrient uptake to regulate enzymes distributed throughout the entire metabolic network. The swift production of activator metabolites after nutrient uptake underscores a coordinated and energy-efficient strategy employed by cells to modulate metabolic pathways. This timely response ensures a rapid and effective adaptation to nutrient availability, enabling cells to generate metabolic flux and efficiently utilize nutrients for diverse cellular processes.

## Methodology

### Reconstructing the cell-intrinsic activation network of *Saccharomyces cerevisiae*

The cell-intrinsic enzyme-metabolite activation-interaction network (‘activation network’) of *Saccharomyces cerevisiae* was reconstructed by combining the activating compounds, collected from the BRaunschweig ENzyme DAtabase (BRENDA) database ^17^, with the *S. cerevisiae* genome-scale metabolic model ^16^. The list of all possible activating compounds for every metabolic enzyme of the *S. cerevisiae* metabolic model (total of 469 unique EC numbers) was downloaded from the BRENDA database using the *getActivatingCompound* function ^17^. In total 2102 compounds, which included both cellular and non-cellular compounds, were reported to activate 313 metabolic enzymes. In addition, several activating compounds appear to use multiple synonyms in the BRENDA database, and this redundancy was removed after curation. A unique ID from the Kyoto Encyclopedia of Genes and Genomes (KEGG) database ^19^ was assigned to each activating compound and iMM904 model compound using a combination of the merging algorithms (http://cts.fiehnlab.ucdavis.edu/) and manually examining the KEGG database synonyms ^19^. Then, all non-cellular compounds were removed from the analysis by mapping the total activating compounds with the list of metabolic compounds of the model. In total 201 metabolites were successfully mapped to iMM904 metabolites, activating 187 enzymes. Finally, the activation network was created by joining enzymes with metabolite nodes when an enzyme is reported to be activated by a metabolite, constituting a total of 874 edges in the network (Supplementary File 1).

### Enzyme and metabolite essentiality prediction

The essentiality of enzymes and metabolites was predicted by performing in-silico knockout experiments on the iMM904 model using Cobra Toolbox ^45^. For an enzyme (or metabolite), both the lower and upper flux bounds of all associated reactions were set to zero. Then the maximum growth was predicted by the Flux Balance Analysis (FBA) approach on a minimal media constraints setting. If the predicted growth in the enzyme (or metabolite) knock-out condition was reduced by more than 90% compared to the normal condition growth rate then the enzyme (or metabolite) was considered essential. The default minimal media setting of the model was used for simulation.

### Enzyme and metabolite classes

For enzyme and metabolite classification, the KEGG database definition was used ^19^. The enzymes were grouped into 6 enzyme classes including oxidoreductases, transferases, hydrolases, lyases, isomerases, ligases, and translocases. For metabolite classification, we used the KEGG database definition. Various small compound groups were merged into one larger group. In total, we use 8 metabolite group nomenclature.

### Biochemical pathway activation

Biochemical pathway information was taken from the KEGG database ^19^. For every pathway, the order of enzymatic reactions was based on KEGG Orthology (KO) - *Saccharomyces cerevisiae* (https://www.genome.jp/brite/sce00001.keg). The enzymes which are not present in the metabolic model reconstruction were removed.

### Network of pathway-path activation interactions

Pathway-pathway activation network was created based on the biochemical pathway definition of the iMM904 metabolic model ^16^. First, for each pair of pathways, the activation frequency was calculated by counting the number of metabolites from one pathway activating the enzymes of another pathway. This frequency was calculated for all pairs of metabolic pathways, and an activation frequency table for pathways was generated. Then, using a hypergeometric statistical test, the significance of activation between each pair of pathways was calculated. Finally, the pathway-pathway activation network was reconstructed by joining two pathways if the P (pathway activating another pathway) < 0.05.

### Transcriptome data analysis and gene co-expression

The raw microarray gene expression data from 207 different studies from the Genome2 chips, covering 3115 samples and 665 unique experimental conditions, were extracted from the ArrayExpress database ^18^ and processed by applying the limma package functions in R ^46^. The RAW cell files were processed, and the average normalized gene expression data were calculated for each unique condition. Using the transcriptome profile, the Pearson correlation test was performed for each pair of genes by the *rcorr* function in R. Adjusted P value was calculated by Benjamini-Hochberg (BH) multiple testing procedure using the *p.adjust* function. Two genes were considered co-expressed if r>0.85 & adjusted P value <0.001.

### Core and variable genes

The core and variable gene information was taken from Peter et al ^47^ which reported the whole-genome sequencing and phenotyping of 1,011 *Saccharomyces cerevisiae* isolates. Out of the total 6081 non-redundant S288C strain ORF, 1144 ORFs are variable and the remaining 4937 are core ORF that are present in all isolates.

### Shortest path length for metabolites and enzymes in the metabolic network

In a graph, the shortest path length is the minimum number of edges between the two nodes. In order to calculate the shortest path length for metabolites, first, the genome-scale model was converted into a graph where nodes represent the metabolite and edges are formed when metabolites are connected by a chemical reaction. Then, for every metabolite, the shortest path length from glucose metabolite was calculated by counting the minimum number of edges between the two metabolites. Similarly, to calculate the shortest path length for enzymes, the iMM904 model was converted into a graph where nodes represent the reactions/enzymes, and edges are established between nodes if a substrate of one reaction is a product of another reaction/enzyme. For each reaction/enzyme, the minimum number of edges from the glucose transport reaction was considered the shortest path length of the reaction/enzyme. The shortest path length for metabolites and enzymes were calculated by *shortest.paths* function from the igraph package in R. Some of the highly connected metabolites including h, h2o, adp, atp, amp, pi, nad, and nadh were removed from creating these graphs.

### Genetic interactions comparison

The genetic interaction data were obtained from Costanzo et al ^48^.

### Computational analysis

Networks were created using functions from the igraph package in R. All the statistical analyses were performed in R. The cobra toolbox was used for simulating genome-scale metabolic reconstruction ^45^. Gene expression data were processed using the limma package in R ^46^.

## Supplementary information

**Supplementary Figure S1:** The distribution of degrees among metabolites and enzymes in the network of enzyme-metabolite activatory interactions follow a power-law distribution.

**Supplementary Figure S2:** The shortest path lengths of activator and non-activator metabolites from all nutrient uptake metabolites indicate that activators exhibit significantly shorter path lengths compared to non-activator metabolites.

**Supplementary File 1:** Enzyme-metabolite activatory interactions network of *Saccharomyces cerevisiae*.

## Funding

SM Al-Zubaidi, MIN and MTA were supported by the UAE University internal research grants (startup grant code G00003688, UPAR grant code G00004152, and SURE+ grant G00004383).

## Author Contributions

The study was conceived by MTA, and analysis was performed by SM Al-Zubaidi, MIN and MTA. MR, RAN and MTA provided critical guidance and interpretation. MTA wrote the first draft of the manuscript, and MR and RAN contributed in writing.

## Competing Interest

The authors declare no competing interest.

